# Comparative Transcriptional Responses of Human Blood to Neutron and Photon Irradiation

**DOI:** 10.64898/2026.08.28.747800

**Authors:** Ahmed Salah, Daniel Wollschläger, Ulrich Giesen, Heinz Schmidberger, Federico Marini, Sebastian Zahnreich

**Author notes:** Correspondence: Sebastian Zahnreich. Federico Marini and Sebastian Zahnreich jointly supervised the work.

## Abstract

Despite the well-known health risks of neutron exposures, key gaps remain in understanding neutron-induced molecular responses and identifying reliable biodosimetric markers that distinguish neutrons from photon exposure. We provide the first genome-wide analysis of the human blood transcriptional response to an accelerator-derived fission-like spectrum of neutrons versus photons, evaluating transcriptomic relative biological effectiveness (RBE) and radiation quality-discriminating gene signatures. Whole blood from healthy donors was irradiated *ex vivo* with X-rays (140 kV, 0-4 Gy, n = 3) or neutrons (0.1-8 MeV, 0-1 Gy, n = 2), incubated for 6 h or 24 h, and processed for RNA sequencing from peripheral blood mononuclear cells (PBMCs). Neutrons were markedly more potent than X-rays at inducing differentially expressed genes (DEGs) at equal doses, showing a peak response 6 h post-irradiation followed by a decline. In contrast, X-rays caused a continuous increase in DEGs up to 24 h (neutrons vs. X-rays at 1 Gy: 1,449 vs. 121 DEGs at 6 h; 996 vs. 621 DEGs at 24 h). A universal p53-centered 34-gene signature, including *FDXR*, *EDA2R*, *GADD45A*, and *ZMAT3*, showed highly monotonic dose responses (Spearman correlation coefficient ≈ 1) across donors, radiation qualities, and timepoints. Additionally, difference-in-differences analysis identified radiation quality-discriminating genes only at 6 h, with transcriptional convergence observed by 24 h, suggesting a very narrow time window for biodosimetric differentiation. We identified a neutron-specific gene signature driven by cGAS-STING-NF-κB signaling (*RELB*, *NFKB1*, *C3*, *MALAT1*) and suppression of B-cell and myeloid identity genes (*IGHD*, *TCL1A*, *CLEC7A*, *TLR2*), defining a biologically coherent neutron quality index with distinct immunomodulatory effects. For the first time, we assessed neutron RBEs at the gene, pathway, and global transcriptomic levels in a human blood model, reporting a global transcriptomic neutron RBE of 1.30 (95% CI: 1.14–1.49) at 6 h and 1.21 (95% CI: 1.14–1.28) at 24 h, providing a valuable basis for biodosimetry in mixed-field exposure scenarios. Our findings advance the mechanistic understanding of neutron radiation responses and support the development of biodosimetric approaches for mixed-field exposure scenarios.

## Introduction

Neutron exposure poses elevated health risks compared with photon radiation due to its energy-dependent increased relative biological effectiveness (RBE). Depending on the biological endpoint, neutron RBEs can reach values of up to 20 at energies of approximately 0.3–0.4 MeV (1). Neutron exposure is relevant in diverse scenarios including nuclear environments (occupational exposure, accidental mass-casualty events), cosmic radiation settings (manned spaceflight), and radiotherapeutic applications (particle therapy, boron neutron capture therapy) (1,2). Despite a historically well-established radiobiological distinctiveness of neutrons, the molecular mechanisms underlying their cellular responses, particularly signaling pathways and transcriptional dynamics, remain poorly characterized (2). Research has historically focused on DNA damage in peripheral blood lymphocytes for biological dosimetry following radiation accidents or occupational exposure. Standard cytogenetic assays, including dicentric chromosome analysis and micronucleus scoring, provide dose estimates but offer limited insight into broader biological effects (3–6).

Beyond their greater DNA-damaging potential and hematotoxicity relative to low linear energy transfer (LET) radiation, complex radiation exposure scenarios involving neutron contributions have been linked to immunomodulatory effects, including systemic inflammation and premature immunosenescence (7–11). We previously demonstrated that *ex vivo* photon irradiation of human whole blood elicits a biphasic transcriptomic response: DNA damage-associated signaling pathways are activated at doses below 1 Gy, whereas robust inflammatory pathway up-regulation emerges only at doses ≥ 2 Gy (12). In contrast, comparable immunologically relevant responses following *ex vivo* neutron irradiation of human blood have been reported only at a relatively high dose of 1 Gy (13), despite their substantially greater RBE for cytogenetic endpoints under the same irradiation conditions, ranging between 4 and 7 (6,14). Whether high-LET neutron exposure induces profound alterations of the peripheral immune landscape, and whether these responses differ mechanistically from those triggered by low-LET photons, remains a critical unresolved question with implications for therapeutic applications, occupational safety, and accidental exposure risk assessment.

Over the past two decades, transcriptomic analyses of the radiation response in peripheral blood have emerged as a valuable complement to biodosimetry, enabling identification of dynamic molecular signatures beyond dose assessment (13,15–19). These approaches can be especially useful for investigating immunological and hematotoxic effects, promoting acute and long-term health outcomes. However, transcriptomic studies of neutron exposure remain very limited. Existing research has primarily examined neutron energy spectra resembling those of the Hiroshima atomic bomb detonation or an improvised nuclear device, using *ex vivo* human or murine whole-blood models or *in vivo* mouse exposures (13,15–19). These studies relied on microarray or qRT-PCR-based analyses. In contrast, evidence is still lacking regarding genome-wide coverage of the transcriptomic responses through comprehensive RNA-sequencing (RNA-seq) analyses that are capable of revealing regulatory networks, global cellular responses, and biomarker candidates. Closing this knowledge gap is critical for distinguishing the biological effects of different radiation qualities through transcriptional signatures and identifying reliable biomarkers of neutron exposure for biodosimetric purposes, but also to improve our understanding of tissue damage and immune modulation.

To address these needs in neutron biodosimetry and health risk assessment, we performed a systematic RNA-seq analysis of transcriptional responses in peripheral blood mononuclear cells (PBMCs) isolated from human blood exposed ex vivo to an accelerator-generated fission-like spectrum of neutrons (0.1-8 MeV) vs. photon irradiation. By linking molecular radiobiology with real-world exposure scenarios, this study provides new insights into neutron-specific cellular and hematologic radiation responses and associated health risks, while supporting advances in radiation protection for nuclear, aerospace, and clinical exposures.

## Methods

### Blood sampling, irradiation, and culturing

Whole blood was collected on the morning of the same day of irradiation from three healthy donors by venipuncture into EDTA tubes (S-Monovette® EDTA K3E, Sarstedt, Nuembrecht, Germany). Ethical approval was obtained from the Ethics Committee of the Rhineland-Palatinate Chamber of Physicians [No. 2023-17191], and all donors provided written informed consent.

X-ray irradiation of blood samples from two male donors and one female donor was conducted using a D3150 X-Ray Therapy System (Gulmay Ltd., Surrey, UK) at 140 kV with a dose rate of 3.6 Gy/min at room temperature as previously described in detail (12). Samples were exposed to 0.1, 0.25, 0.5, 1, 2, and 4 Gy, with sham-irradiated (0 Gy) controls maintained under identical conditions in the control room.

Neutron irradiation of blood samples from the same two male donors as for X-ray irradiation, enabling cross-radiation comparisons while controlling for inter-donor variability, was performed at the Physikalisch-Technische Bundesanstalt (PTB) accelerator facility (PIAF) in Braunschweig, Germany, using a 2 MV Tandem accelerator. The neutron field was generated by a neutron reaction induced by a 3.4 MeV deuteron beam with currents up to 50 µA on a 4 mm-thick, water-cooled beryllium disc target. The resulting neutron energy spectrum spans from very low energies up to approximately 8 MeV, with a tissue-kerma-weighted mean energy of approximately 2.5 MeV, representative of a fission-like spectrum. Blood samples were positioned on a rotating device at 585 mm from the target and approximately 180 mm from the outer edge of the collimator system, at room temperature (20.0 ± 1.0° C). The absorbed dose to tissue was determined according to ICRU Report No. 45 (20) using a tissue-equivalent A-150 ionization chamber (EXRADIN, T2-#381, Standard Imaging, Inc.) filled with tissue-equivalent gas, calibrated in the ⁶⁰Co reference field of PTB. The photon contamination component of the field was determined to be (10 ± 2)% of the total dose. Beam charge on the beryllium target served as the irradiation monitor. Samples were exposed to nominal doses of 0, 0.1, 0.25, 0.5, and 1.0 Gy, corresponding to absorbed doses to tissue of 0.098, 0.245, 0.490, and 0.981 Gy, respectively, each with a measurement uncertainty of 5.7% and a dose rate of approximately 0.019-0.020 Gy/min.

### RNA isolation from irradiated blood samples

After X-ray or neutron irradiation, blood samples were diluted 1:1 with pre-warmed (37 °C) X-VIVO™ 15 medium (Lonza Group Ltd., Basel, Switzerland), transferred into cell suspension flasks and incubated at 37 °C with 5% CO₂ in a humidified atmosphere for 6 h or 24 h. Following incubation, PBMCs were isolated by density gradient centrifugation. Two volumes of Erythrocyte Lysis Buffer (Qiagen) were added to the extracted PBMCs and incubated on ice for 5-10 min to remove the remaining red blood cells, followed by centrifugation at 400 × g for 10 min at 4° C. The supernatant was discarded, and the cell pellet was washed with 10 ml PBS to remove residual buffer. Total RNA was then extracted from the cell pellet using the NucleoSpin® RNA Plus kit (Macherey-Nagel) according to the manufacturer’s protocol.

### RNA sequencing

RNA library preparation and transcriptome sequencing were performed by Novogene GmbH (Munich, Germany) according to the company’s protocols. Libraries were prepared using the Novogene NGS RNA Library Prep Set (PT042) and quantified using a Qubit 2.0 fluorometer and real-time PCR. Size distribution was assessed on an Agilent 2100 Bioanalyzer. Sequencing was performed on a NovaSeq X platform, generating paired-end 150 nt reads. Base calling was performed using CASAVA, and data was converted to FASTQ format using bcl2fastq (v2.20.0.422).

### Processing of RNA-seq data

#### Quality control, exploratory data analysis, and Differential gene expression analysis

Raw sequencing data quality was assessed using FastQC (v0.12.1; https://www.bioinformatics.babraham.ac.uk/projects/fastqc/). Transcript abundance was estimated using Salmon (21) (v1.10.3) with a decoy-aware transcriptome index based on GENCODE release 49, and summarised to gene level using the tximeta R package (22) (v1.28.3).

Principal component analysis (PCA) was performed using variance-stabilized transformation (VST)-normalized gene expression of the 500 most variable genes.

Differential expression analysis was performed using DESeq2 (23) (v1.50.2) and limma (24) (v3.66.0), with a false discovery rate (FDR) threshold of 0.05. Sham-irradiated (0 Gy) samples for both radiation qualities served as the reference level. The statistical model accounted for dose effects and inter-donor variability. Log2 fold change (LFC) effect sizes were estimated using the apeglm shrinkage estimator (25) (v1.32.0). To compare radiation responses at 6 h and 24 h post-irradiation, a time-series analysis was implemented using DESeq2 by specifying dose × time interaction terms. Neutron 0.25 Gy 24 h samples were excluded due to insufficient yield and quality.

To identify genes with differential dose-response slopes between radiation types, a difference-in-differences (DiD) design was implemented in limma-voom. The DiD contrast was defined as:

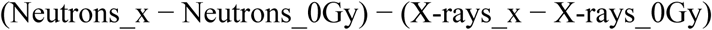

where subscripts denote irradiated (x Gy) and 0 Gy baseline (0) conditions. Seven contrasts were computed across shared doses and time points.

Temporal dynamics were characterized using a baseline-corrected time point DiD, also implemented in limma-voom:

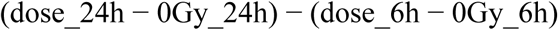

This formulation removes culture-drift artefacts independently at each time point. Genes were considered significantly regulated if adjusted p-value (p_adj_) ≤ 0.05.

### Meta-correlation analysis

To quantify within-donor dose-response relationships and identify genes showing consistent monotonic responses to radiation across donors and experimental conditions, a within-donor Spearman meta-correlation analysis was performed (26). For each gene and experimental setting (defined as radiation type × timepoint: neutrons 6 h, neutrons 24 h, X-rays 6 h, X-rays 24 h), Spearman correlation coefficients between radiation dose (0–1 Gy) and VST-normalised expression were computed independently for each donor, then pooled across donors using Fisher z-transformation meta-analysis as implemented in the metacor function of the R meta package (27) (v8.5.0) (using the common-effect model), with results back-transformed to the correlation scale via the hyperbolic tangent function. FDR correction was applied per arm using the Benjamini-Hochberg method. Inter-donor consistency was assessed by computing the standard deviation of per-donor Spearman r values across arms for each gene; genes where all donors showed the same direction of correlation in every experimental setting and a mean inter-donor SD ≤ 0.10 were considered robustly consistent across individuals. Universal dose-response biomarker candidates were pre-selected from genes showing FDR-significant differential expression in at least two contrasts within each of the experimental groups, restricted to doses ≤ 1 Gy to ensure consistency with the meta-correlation dose range. Best-performing genes were selected based on the lower bound of their absolute pooled Spearman r across all experimental groups, ensuring consistent dose-response (|r| at least 0.8, FDR < 0.05 in all four arms) across all conditions rather than relying on average performance.

### Pathway enrichment analysis

Over-representation analysis (ORA) of differentially expressed genes (DEGs) was performed using clusterProfiler (28,29) (v4.18.4) with the enrichGO and gene set enrichment analysis (GSEA) functions, using all expressed genes as the background. Enrichment results were visualised using the GeneTonic package (30) (v3.4.0). Gene expression profiles were displayed as heatmaps of standardized Z-scores derived from VST values.

### Transcriptomic relative biological effectiveness

#### Global RBE

A shared radiation-responsive gene signature was derived by identifying genes significantly differentially expressed in both radiation arms at each time point. The top 50 upregulated and 50 downregulated genes, ranked by mean absolute LFC, were selected. VST expression values were Z-scored per gene, direction-weighted (1 for upregulated, −1 for downregulated), and averaged per sample to yield a composite radiation score. Donor-level linear regression of score ∼ dose was performed within the shared 0-1 Gy range, and the global RBE was calculated as slope (neutrons) / slope (X-rays). Confidence intervals were estimated by donor-level bootstrap resampling (n = 2,000 iterations).

#### Gene-level RBE

A DESeq2 model was fitted to the 48 shared-dose samples (0-1 Gy). Neutron and X-ray dose-response effects were extracted via explicit group contrasts with ashr shrinkage (31):

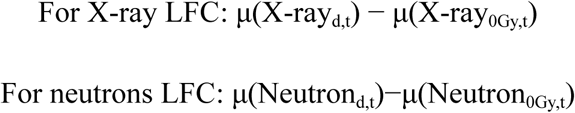

Where d denotes dose (0, 0.1, 0.25, 0.5, or 1 Gy) and t denotes timepoint (6h or 24h), and μ denotes the group mean log2 expression estimated from the DESeq2 model.

Gene-level RBE was computed exclusively for genes with a measurable dose-dependent response in both radiation arms, as genes with absent or near-zero responses in one experimental setting will produce unstable or biologically uninterpretable RBE ratios regardless of significance in the other experimental setting. Gene-level RBE was calculated as neutron LFC / X-ray LFC at each dose × time point combination, subject to a four-step filter cascade: (F1) model convergence and absence of dispersion outliers; (F2) baseMean ≥ 5; (F3) |X-ray LFC| ≥ 0.5 to stabilize the denominator; (F4) interaction p_adj < 0.05 or both arms p_adj_ < 0.05.

#### Pathway-level RBE

Gene Set Variation Analysis (GSVA) was performed using the GSVA package (v2.4.9) (32). Gene Ontology (GO) annotations were obtained from the org.Hs.eg.db package (v3.22.0) and Hallmark gene sets from MSigDB. GSVA was applied to the top 5,000 high-variance genes from the donor-corrected VST matrix, using GO Biological Process (GO:BP) gene sets (10-500 genes) and MSigDB Hallmark gene sets separately. Per-sample pathway enrichment scores were modelled using limma linear models, fitted separately for each radiation type and time point within the shared 0-1 Gy range. The dose-response slope coefficient was taken as pathway activity per Gy. Pathway RBE was calculated as slope (neutron) / slope (X-ray), after applying filters for X-ray statistical significance (p_adj_ < 0.05), concordant slope direction, and denominator stability (|X-ray slope| ≥ 0.05 for GO:BP; ≥ 0.10 for Hallmarks).

## Results

Blood samples from three and two healthy donors were irradiated with X-rays (0-4 Gy) or neutrons (0-1 Gy), respectively, and incubated for 6 or 24 h to characterize and compare transcriptional dose-response relationships and kinetics across radiation qualities.

### Time post-exposure is the dominant driver of transcriptional variation

PCA (Figure 1A) showed that PC1, explaining 49.7% of the variance, separated samples by post-exposure time, whereas PC2, explaining 9.5%, distinguished samples by radiation quality. This pattern indicates that post-exposure time is the primary driver of transcriptional variation, followed by radiation quality, while donor-specific effects were effectively controlled through the study design and statistical modeling.

**Figure 1:**
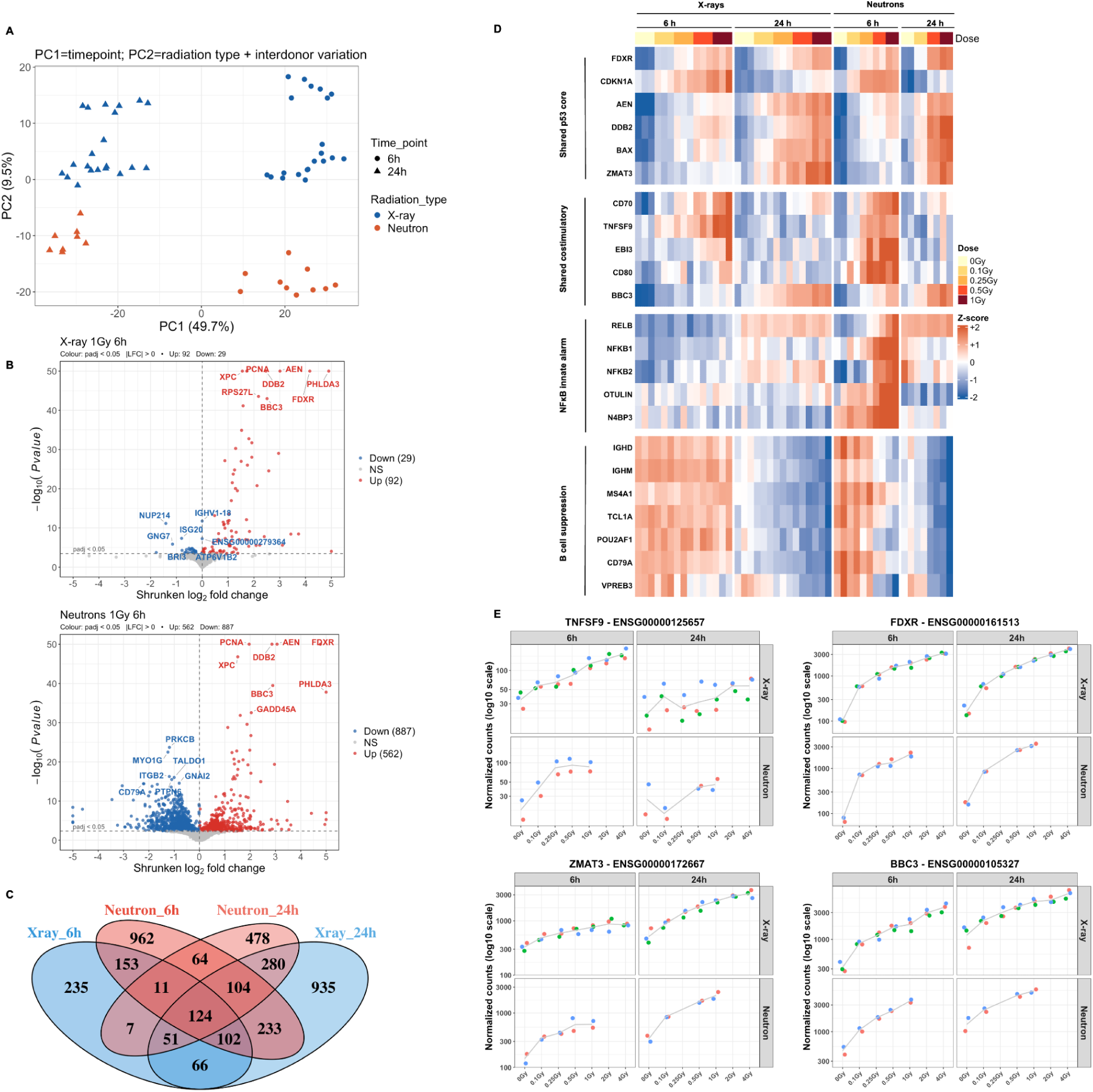
Transcriptional response of peripheral blood mononuclear cells to X-rays and neutrons. (A) Principal component (PC) analysis plot. (B) Volcano plots of differentially expressed genes (DEGs) at 1 Gy 6 h for X-rays (top) and neutrons (bottom). Red dots: significantly upregulated (p_adj_ < 0.05, LFC > 0); blue dots: significantly downregulated; grey dots: non-significant. (C) Four-way Venn diagram showing overlap of DEGs across X-rays and neutrons after 6 or 24 h, aggregated across all shared doses (0–1 Gy). (D) Heatmap showing X-ray and neutron responses for selected representative genes across four functional gene layers: shared p53 core, shared costimulatory alarm, neutron-specific NF-κB innate alarm, and neutron-specific B-cell suppression programme. Columns represent donor samples grouped by radiation quality, time post-exposure, and dose. Colour scale represents Z-score of normalised counts. (E) Dose-response plots for four representative biomarker genes (*TNFSF9*, *FDXR*, *ZMAT3*, *BBC3*) showing normalised counts on a log10 scale across all doses and timepoints after X-rays (top) and neutrons (bottom). Coloured dots represent individual donors.

### Neutrons and X-rays trigger shared and distinct transcriptional programs

Neutrons and X-rays induced dose-dependent transcriptional responses, with neutrons triggering substantially more DEGs than X-rays at equivalent doses across the 0.1–1 Gy range. At 6 h after 1 Gy, neutrons induced 1,449 DEGs compared with 121 for X-rays (Figure 1B).

Both radiation qualities regulated a set of shared and radiation-unique gene expression profiles (Figure 1C). Both radiation qualities activated a universal p53-centered damage response comprising 11 genes significantly upregulated across all doses and timepoints: *FDXR*, *BBC3*, *GADD45A*, *DDB2*, *PHPT1*, *PCNA*, *AEN*, *BAX*, *XPC*, *TRIAP1*, and *PHLDA3*. A transient early costimulatory response comprising 23 shared genes was significantly regulated at 6 h but returned to baseline by 24 h; at 1 Gy, this included *EBI3*, *CD80*, *CD83*, *NFKB2*, *SGPP2*, *MTHFD1L*, and *IGFBP4*. At 24 h, 22, 121, and 295 DEGs were shared between the two radiation types at 0.1, 0.5, and 1 Gy, respectively, including *DDB2* and *ZMAT3* at 0.1 Gy, *TLR6* and *IGKV1-5* at 0.5 Gy, and *MDM2* and *CD22* at 1 Gy (Figure 1D).

The two radiation types showed opposing temporal dynamics. X-ray responses expanded: from 26, 46, 66, 121, 354, and 630 DEGs at 6 h, to 46, 113, 268, 621, 982, and 1,577 DEGs at 24 h (0.1, 0.25, 0.5, 1, 2, and 4 Gy, respectively), whereas neutron responses contracted: from 36, 81, 799, and 1,449 DEGs at 6 h (0.1, 0.25, 0.5, 1 Gy) to 82, 401, and 996 at 24 h (0.1, 0.5, 1 Gy) (Supplementary Table 1).

At low X-ray doses (0.1–0.25 Gy), the 6 h response was largely restricted to canonical p53 targets (*FDXR*, *CDKN1A*, *MDM2*, *BBC3*), broadening at 1–4 Gy to include immune (*CD83*, *CD79A*, *CD180*) and DNA repair genes (*PCNA*, *POLH*). Neutrons activated p53 targets similarly but additionally induced immune-related changes (upregulation of *RELB*, *NFKB1*, *C3*, *CCL22*; downregulation of *VPREB3*, *IGHD*, *NIBAN3*, *CLEC7A*) and the non-coding transcripts *MIR155HG* and *MIAT* already at 0.5–1 Gy.

Each radiation type exhibited a sustained set of genes significantly regulated at both timepoints: 64 genes for neutrons (most strongly: *SECTM1*, *CD300E*, *TNS3*, *PTGDS*) and 66 genes for X-rays. In both signatures, several genes reversed direction over time. In the neutron signature, *M6PR* and *GMFG* were downregulated at 6 h but upregulated at 24 h, whereas *AFF2* and *ZBTB1* showed the opposite pattern. In the X-ray signature, *ARHGAP4* switched from down to up, whereas *ICAM1*, *ABCC4*, *MARS1*, and *CTNND1* switched from up to down.

Among the most strongly induced genes at 1 Gy, 6 h were *EDA2R*, *FDXR*, *EBI3*, *BBC3*, *ZMAT3*, *CD70*, and *TNFSF9* (Figure 1E).

### Identification of robust universal dose-response biomarkers across radiation type and donor background

We first identified broadly applicable candidate biomarkers for biodosimetry across radiation types, doses, time points and donors using a within-donor Spearman meta-correlation analysis to assess dose-response relationships across the shared dose range (0-1 Gy). Candidate biomarkers were pre-selected by requiring significance in at least two differential expression contrasts within each of the four experimental settings, restricted to doses ≤ 1 Gy to ensure consistency with the meta-correlation range, yielding 40 candidate genes. From these, meta-correlation identified 36 strong biomarkers (minimum |r| ≥ 0.8, FDR < 0.05 in all four experimental settings), comprising 35 upregulated genes and one downregulated gene (*ISG20*). 25 genes showed mean and minimum pooled |r| = 1.0, indicating a perfect monotonic dose-response across all donors and experimental settings. This set was strongly enriched for p53 targets and DNA damage response genes, including *GADD45A*, *PCNA*, *XPC*, *FDXR*, *TRIAP1*, *AEN*, *BBC3*, *DDB2*, *POLH*, *ZMAT3*, *PHLDA3*, *FAS*, *TNFRSF10B*, *PPM1D*, *PHPT1*, *BAX*, and *CDKN1A*. Additional genes included cell-cycle regulators (*CCNG1*), apoptosis mediators (*ACTA2*, *CD70*, *GLS2*), and less-characterized radiation response genes (*ZNF79*, *CMBL*, *TMEM30A*, *DCP1B*, *ASTN2*, *PVT1*, *EDA2R*, *MAP4K4*, *ASCC3*, *APOBEC3C*, *FBXO22*). The single downregulated Tier 1 gene, *ISG20*, showed consistent downregulation across all four experimental groups (r = −1.0 for neutrons; r = −0.9 to −1.0 for X-rays).

Of the 36 genes, 34 passed the stringent inter-donor consistency filter (uniform direction across donors, mean donor SD ≤ 0.10, minimum pooled |r| > 0.8), including *BBC3*, *PRKAB1*, and *ASCC3* (Table 1; Figure 2). The highest-confidence radiation biodosimetry candidates were seven genes with mean donor SD = 0 across all four experimental settings, comprising *GADD45A*, *PCNA*, *XPC*, *FDXR*, *TRIAP1*, *AEN*, and *ZNF79*, showing identical dose-response correlations across all donors and conditions. An additional nine genes, including *BBC3*, *PRKAB1*, *ASCC3*, *EDA2R*, *IER5*, *POLH*, *PPM1D*, *PVT1*, and *TNFSF8*, exhibited near-identical responses across donors (mean SD ≤ 0.02).

**Figure 2:**
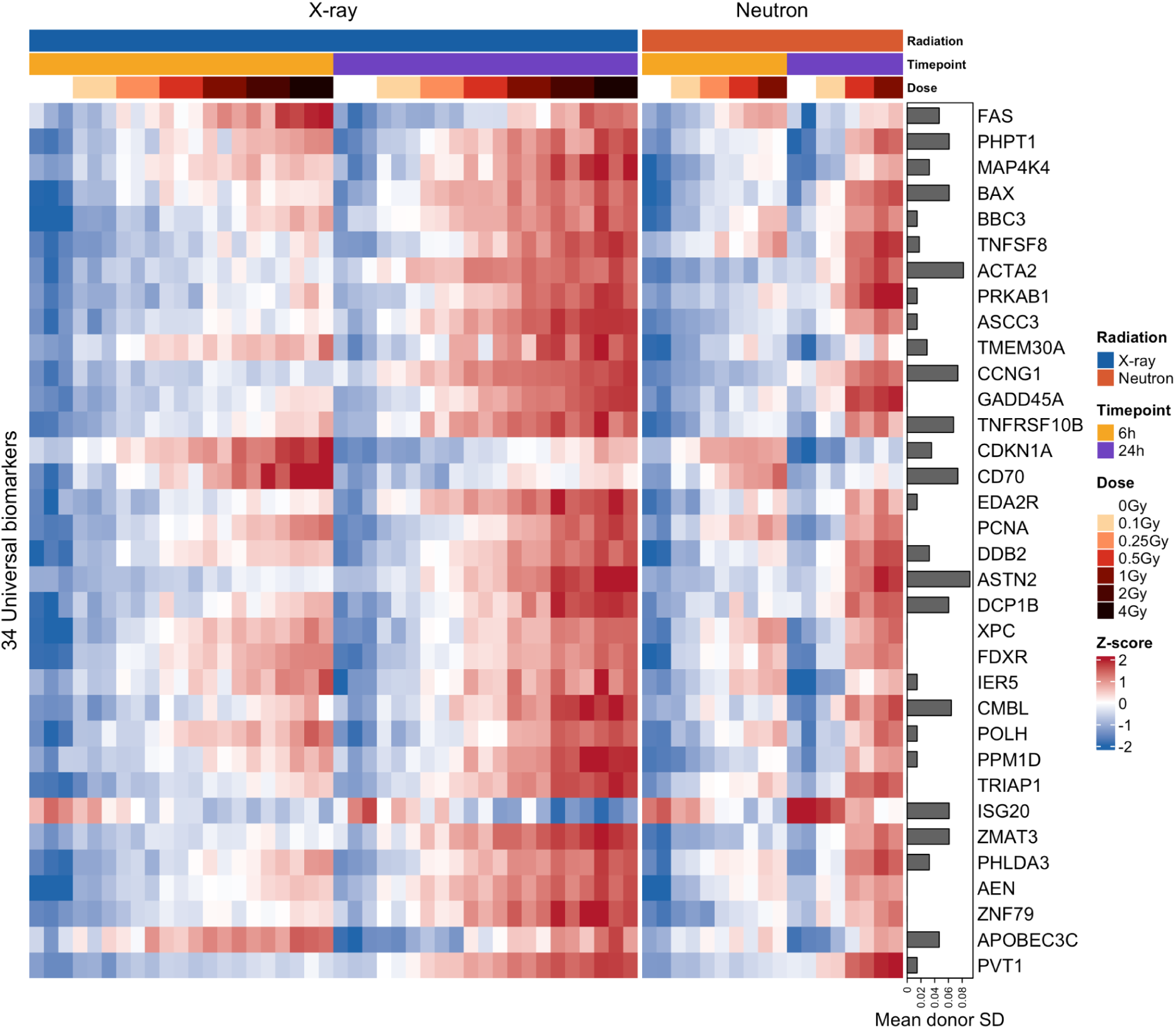
Heatmap of 34 universal biomarker genes showing consistent monotonic dose-response (minimum |r| > 0.80, mean donor SD ≤ 0.10) across all donors, doses, timepoints, and both radiation qualities. The grey bar plot on the right shows mean donor standard deviation for each gene. The colour scale represents the Z-score of normalised counts.

**Table 1:**
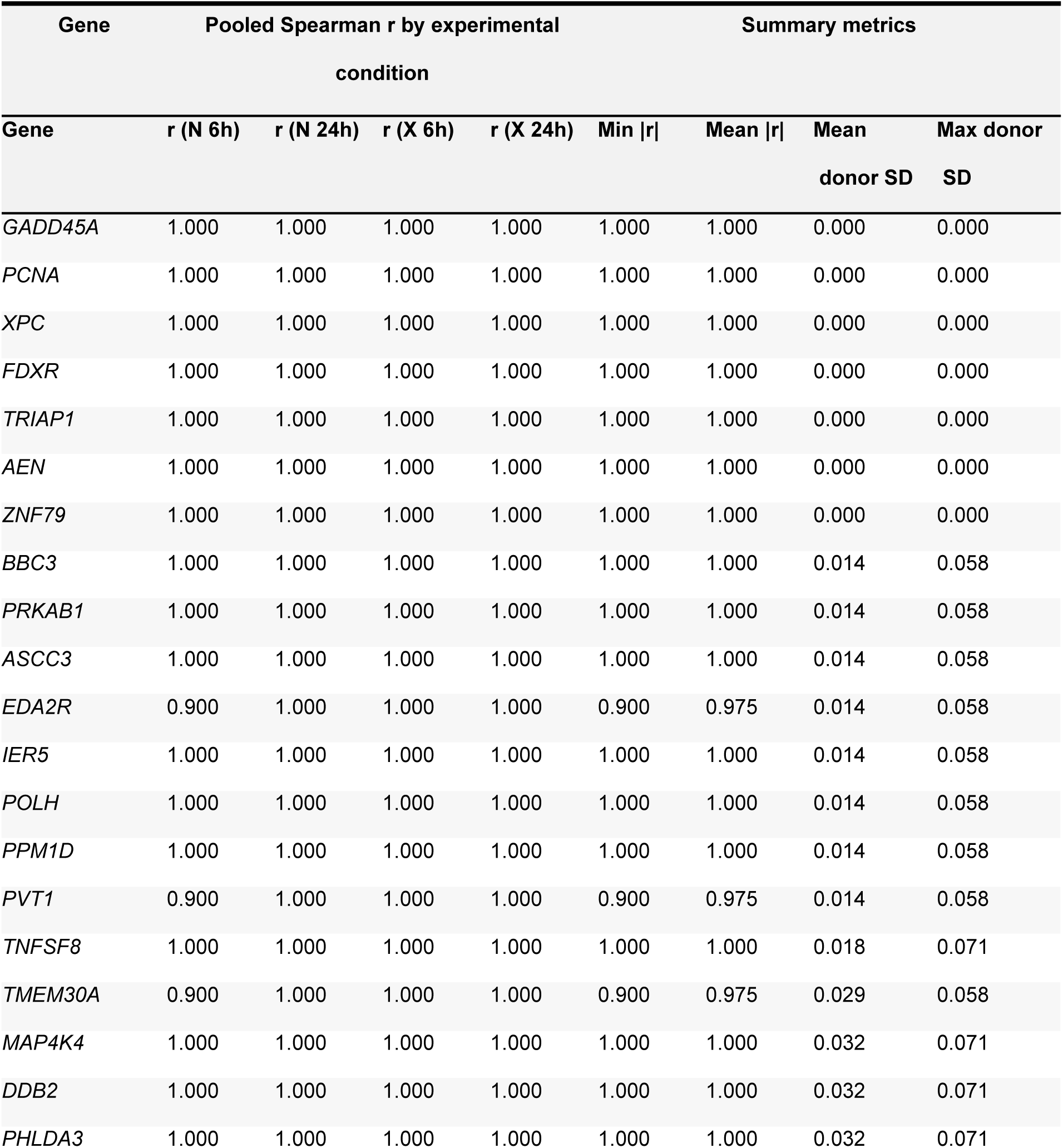
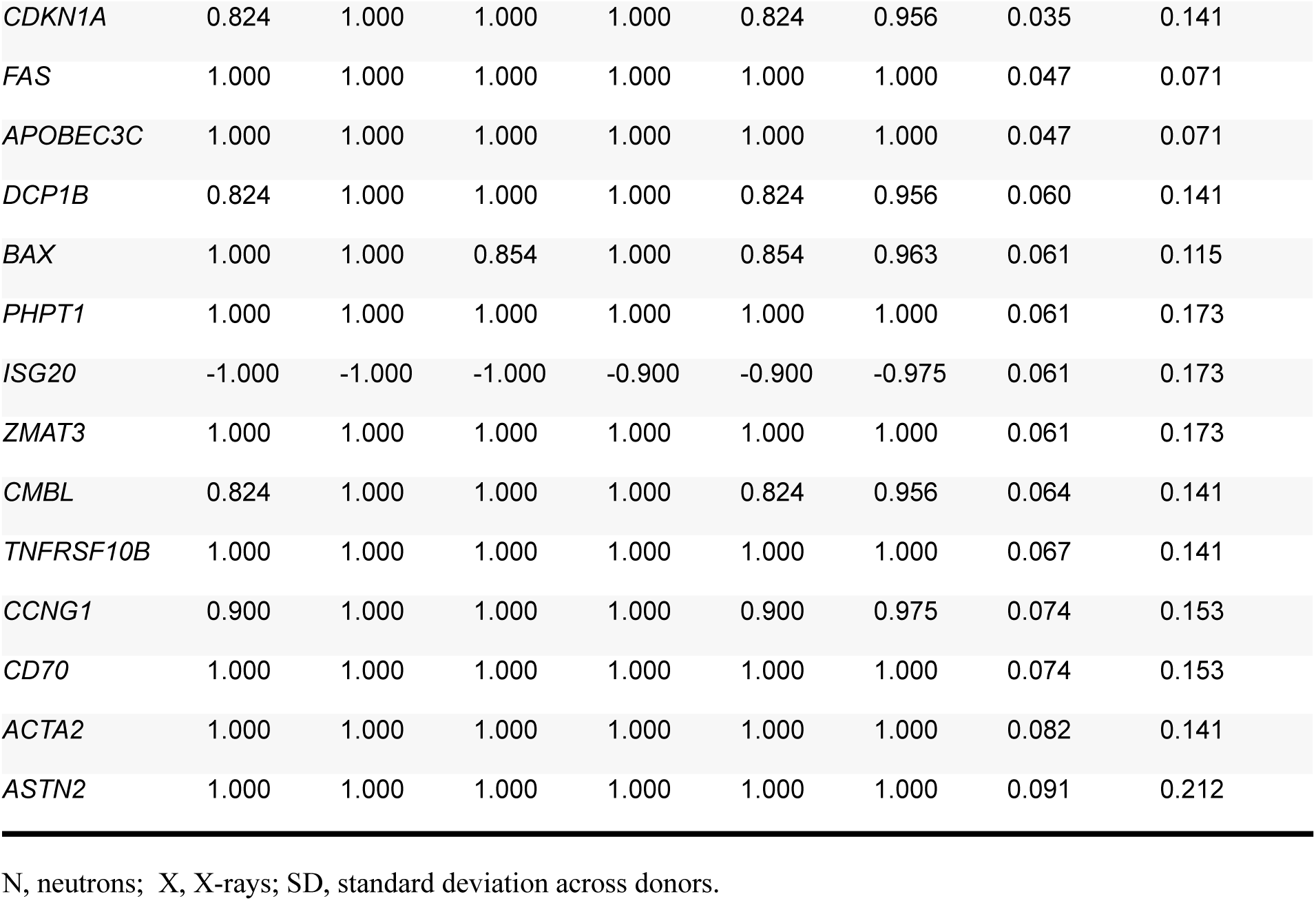
Universal dose-response biomarker genes identified by within-donor Spearman meta-correlation, meeting all three selection criteria (all donors show same direction, mean donor standard deviation ≤ 0.10, min pooled |r| > 0.80) across both radiation qualities and timepoints (6 and 24 h post-irradiation).

| Gene | Pooled Spearman r by experimental condition |  |  |  | Summary metrics |  |  |  |
| --- | --- | --- | --- | --- | --- | --- | --- | --- |
| Gene | r (N 6h) | r (N 24h) | r (X 6h) | r (X 24h) | Min \|r\| | Mean \|r\| | Mean donor SD | Max donor SD |
| *GADD45A* | 1.000 | 1.000 | 1.000 | 1.000 | 1.000 | 1.000 | 0.000 | 0.000 |
| *PCNA* | 1.000 | 1.000 | 1.000 | 1.000 | 1.000 | 1.000 | 0.000 | 0.000 |
| *XPC* | 1.000 | 1.000 | 1.000 | 1.000 | 1.000 | 1.000 | 0.000 | 0.000 |
| *FDXR* | 1.000 | 1.000 | 1.000 | 1.000 | 1.000 | 1.000 | 0.000 | 0.000 |
| *TRIAP1* | 1.000 | 1.000 | 1.000 | 1.000 | 1.000 | 1.000 | 0.000 | 0.000 |
| *AEN* | 1.000 | 1.000 | 1.000 | 1.000 | 1.000 | 1.000 | 0.000 | 0.000 |
| *ZNF79* | 1.000 | 1.000 | 1.000 | 1.000 | 1.000 | 1.000 | 0.000 | 0.000 |
| *BBC3* | 1.000 | 1.000 | 1.000 | 1.000 | 1.000 | 1.000 | 0.014 | 0.058 |
| *PRKAB1* | 1.000 | 1.000 | 1.000 | 1.000 | 1.000 | 1.000 | 0.014 | 0.058 |
| *ASCC3* | 1.000 | 1.000 | 1.000 | 1.000 | 1.000 | 1.000 | 0.014 | 0.058 |
| *EDA2R* | 0.900 | 1.000 | 1.000 | 1.000 | 0.900 | 0.975 | 0.014 | 0.058 |
| *IER5* | 1.000 | 1.000 | 1.000 | 1.000 | 1.000 | 1.000 | 0.014 | 0.058 |
| *POLH* | 1.000 | 1.000 | 1.000 | 1.000 | 1.000 | 1.000 | 0.014 | 0.058 |
| *PPM1D* | 1.000 | 1.000 | 1.000 | 1.000 | 1.000 | 1.000 | 0.014 | 0.058 |
| *PVT1* | 0.900 | 1.000 | 1.000 | 1.000 | 0.900 | 0.975 | 0.014 | 0.058 |
| *TNFSF8* | 1.000 | 1.000 | 1.000 | 1.000 | 1.000 | 1.000 | 0.018 | 0.071 |
| *TMEM30A* | 0.900 | 1.000 | 1.000 | 1.000 | 0.900 | 0.975 | 0.029 | 0.058 |
| *MAP4K4* | 1.000 | 1.000 | 1.000 | 1.000 | 1.000 | 1.000 | 0.032 | 0.071 |
| *DDB2* | 1.000 | 1.000 | 1.000 | 1.000 | 1.000 | 1.000 | 0.032 | 0.071 |
| *PHLDA3* | 1.000 | 1.000 | 1.000 | 1.000 | 1.000 | 1.000 | 0.032 | 0.071 |
| *Blood Transcriptomics After Neutrons vs. Photons* |  |  |  |  |  |  |  |  |
| *CDKN1A* | 0.824 | 1.000 | 1.000 | 1.000 | 0.824 | 0.956 | 0.035 | 0.141 |
| *FAS* | 1.000 | 1.000 | 1.000 | 1.000 | 1.000 | 1.000 | 0.047 | 0.071 |
| *APOBEC3C* | 1.000 | 1.000 | 1.000 | 1.000 | 1.000 | 1.000 | 0.047 | 0.071 |
| *DCP1B* | 0.824 | 1.000 | 1.000 | 1.000 | 0.824 | 0.956 | 0.060 | 0.141 |
| *BAX* | 1.000 | 1.000 | 0.854 | 1.000 | 0.854 | 0.963 | 0.061 | 0.115 |
| *PHPT1* | 1.000 | 1.000 | 1.000 | 1.000 | 1.000 | 1.000 | 0.061 | 0.173 |
| *ISG20* | -1.000 | -1.000 | -1.000 | -0.900 | -0.900 | -0.975 | 0.061 | 0.173 |
| *ZMAT3* | 1.000 | 1.000 | 1.000 | 1.000 | 1.000 | 1.000 | 0.061 | 0.173 |
| *CMBL* | 0.824 | 1.000 | 1.000 | 1.000 | 0.824 | 0.956 | 0.064 | 0.141 |
| *TNFRSF10B* | 1.000 | 1.000 | 1.000 | 1.000 | 1.000 | 1.000 | 0.067 | 0.141 |
| *CCNG1* | 0.900 | 1.000 | 1.000 | 1.000 | 0.900 | 0.975 | 0.074 | 0.153 |
| *CD70* | 1.000 | 1.000 | 1.000 | 1.000 | 1.000 | 1.000 | 0.074 | 0.153 |
| *ACTA2* | 1.000 | 1.000 | 1.000 | 1.000 | 1.000 | 1.000 | 0.082 | 0.141 |
| *ASTN2* | 1.000 | 1.000 | 1.000 | 1.000 | 1.000 | 1.000 | 0.091 | 0.212 |

N, neutrons; X, X-rays; SD, standard deviation across donors.

### Pathway enrichment reveals conserved p53 core and dose-dependent immune modulation

ORA confirmed the gene-level findings (Supplementary Table 2). The p53 Pathway Hallmark was the most significantly enriched term in both radiation types, appearing at every dose and timepoint, from 0.1 Gy (dominant) to 4 Gy (top term). Apoptosis and DNA repair were consistently co-enriched with p53, supporting a common p53-mediated stress response (Figure 3A and C). GSEA further reinforced these findings (Supplementary Table 3; Supplementary Figure 1). The p53 pathway showed the highest positive normalized enrichment score (NES, 2.58-3.12) across essentially all contrasts. GO-enriched terms yielded similar results, with intrinsic apoptotic signaling, p53 signal transduction, and cellular responses to radiation among the top enriched pathways from the lowest neutron and X-ray doses onward. Additionally, ubiquinone biosynthesis and metabolic processes were consistently positively enriched across all experimental contrasts (GO:BP NES 2.07-2.41). Notably, the enrichment patterns of suppressive immune effector programs that persisted up to 24 h post-exposure diverged between radiation qualities in a dose-dependent manner.

**Figure 3:**
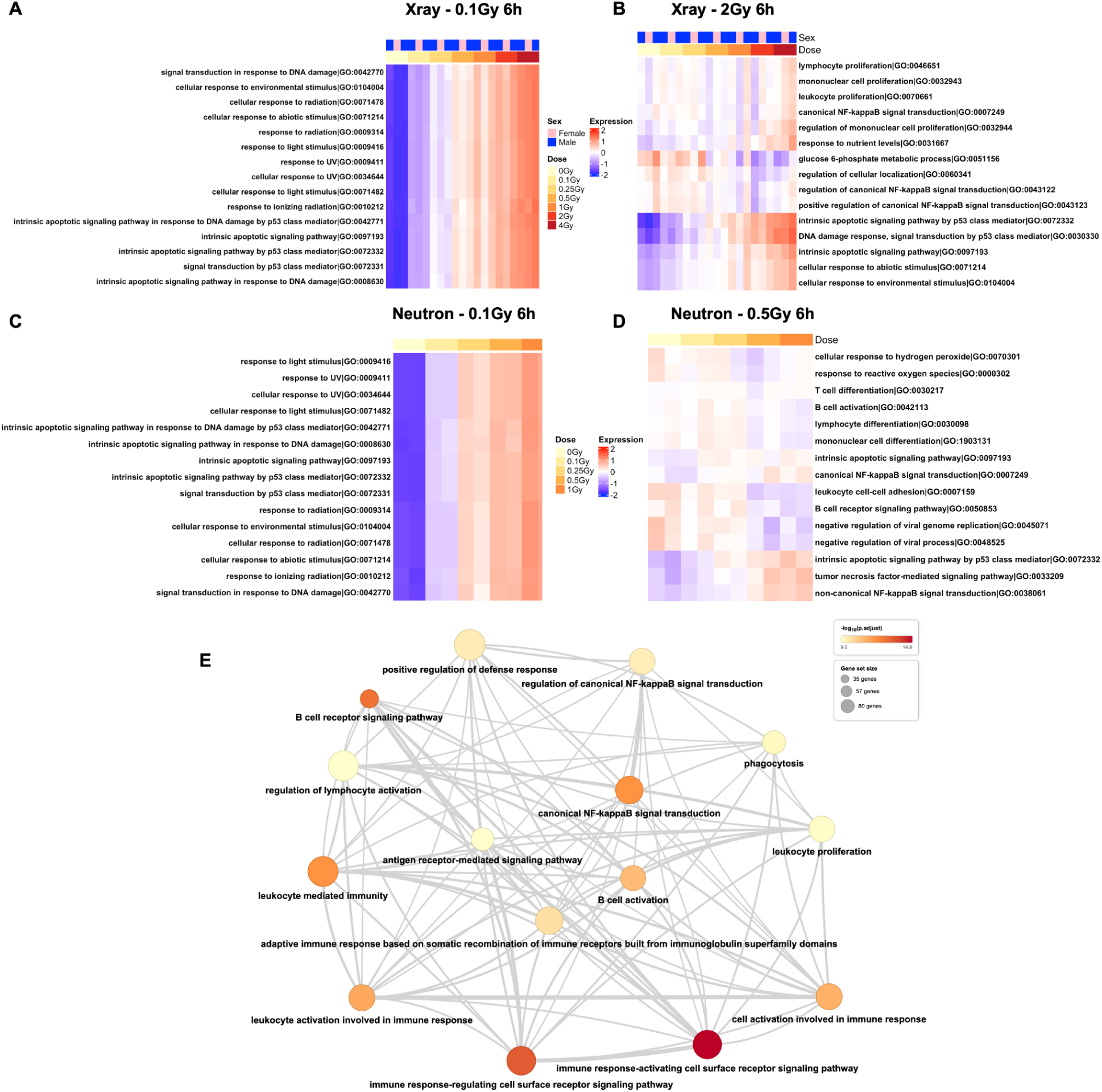
Pathway enrichment heatmaps 6 h after (A) 0.1 Gy X-rays, (B) 2 Gy X-rays, (C) 0.1 Gy neutrons and (D) 0.5 Gy neutrons. Columns represent individual samples; colour scale indicates enrichment score. (E) Pathway network plot of significantly enriched pathways 6 h after 1 Gy neutrons. Node size represents gene set size; node colour represents −log10(p_adj_).

For X-rays, immune modulation was first detectable 6 h after exposure to 1 Gy and was characterized by suppression of T-cell and lymphocyte proliferation together with activation of TNF-α signaling via NF-κB (Figure 3B). By 24 h, following doses ≥ 2 Gy, the response had expanded to encompass suppression of the innate immune response, Toll-like receptor signaling, the response to tumor necrosis factor, and regulation of type I interferon production. Concurrently, B-cell identity and adaptive immune pathways became increasingly downregulated.

Neutron irradiation elicited a robust immunosuppressive response at substantially lower doses than X-rays (Figure 3D). 6 h after 0.5 Gy, GSEA revealed positive enrichment of TNF-α Signaling and the p53 pathway, together with negative enrichment of Complement, Interferon alpha and Gamma Responses, Allograft Rejection, indicating broader immune-related perturbations than observed after equal X-ray doses. At 1 Gy, affected pathways extended to leukocyte-mediated immunity, humoral immune response, B-cell differentiation, regulation of T-cell activation, and response to type II interferon, confirming a broad immune suppression phenotype driven by neutrons at substantially lower doses compared with X-rays (Figure 3E).

### Radiation quality difference-in-differences analysis identifies early and dose-threshold-dependent divergence

To formally identify genes whose dose-response differs between radiation types, a DiD analysis was implemented across a range of five shared doses (0-1Gy) × timepoint contrasts (Supplementary Table 4). All DiD-significant genes were identified exclusively at 6 h post-exposure, with 280, 19, and only 3 genes detected after 1 Gy, 0.5 Gy, and < 0.5 Gy, respectively. These findings indicate that discrimination between radiation qualities is restricted to an early, dose-dependent response, consistent with the broader DEG landscape.

All DiD-significant genes were classified by cross-referencing LFC and significance within each experimental setting, as the DiD LFC sign alone was insufficient to determine the direction of regulation. The dominant class comprised genes significantly suppressed only after neutrons, with non-significant changes after X-rays, including 184 and 10 genes at 1 Gy and 0.5 Gy, respectively. Representative genes included the B-cell identity markers *VPREB3*, *IGHD*, *TCL1A*, *CD79A*, and *CD163*, as well as *ZNF467*, *FUCA1*, *FAM111B*, and *RASSF2*. The corresponding neutron-specific upregulated class comprised 34 genes at 1 Gy and 6 genes at 0.5 Gy, corresponding to 37 unique DEGs due to overlap between the two doses.

Four genes showed concordant responses across both radiation types with a stronger neutron effect, all at 1 Gy: *NFKB2* showed greater upregulation, whereas *ITGB2*, *TALDO1*, and *GNAI2* displayed more pronounced downregulation. *FCER2* was the only gene showing X-ray-specific induction, and no genes were classified as X-ray-specific downregulated.

### Neutron-preferential activation: NF-κB innate alarm and cGAS-STING amplification

Among the genes activated preferentially by neutrons within the shared 0-1 Gy dose range, a coherent NF-κB signalling pathway was identified 6 h after 0.5 Gy and 1 Gy. Core NF-κB subunits *RELB* (non-canonical) and *NFKB1* (canonical) were activated at both doses alongside *NFKB2*, *TRAF3*, and *OTULIN*. Multiple negative feedback regulators were co-induced, *including NFKBIA*, *NFKBIB*, *TNIP1*, and *BIRC2*. Complement components *C3* and *MALAT1* were among the most strongly induced genes.

Cross-referencing with X-ray responses after 6 h across the full dose range (0-4 Gy) revealed that 24 genes showed no significant X-ray response at 6 h at any tested dose, even at higher doses ≥ 2 Gy, and were designated Class A (X-ray-insensitive at 6 h; Figure 4A). This was the subset with the strongest claim to radiation-quality specificity within the DiD analytical framework. Three mechanistic threads ran through the Class A up genes: (i) cGAS-STING and mitochondrial innate sensing amplification (*PGAM5*, *TSPOAP1*, *ITPR2*, and *N4BP3*); (ii) NF-κB transcriptional program (*CDK5R1*, *PPP1R2*, *SMAD7*, *MED13L*, and *MALAT1*, as the most strongly induced Class A gene, reinforces NF-κB output at the post-transcriptional level); (iii) a potential mechanistic link to B-cell suppression (*BCOR*, *S1PR2*, and *KLHL6*). *C3*, as the most strongly induced Class A gene at 0.5 Gy, represents a central hub of all three complement pathways and a direct NF-κB transcriptional target. Its neutron specificity at 6 h highlighted it as a potential high-LET-specific marker of complement activation.

**Figure 4:**
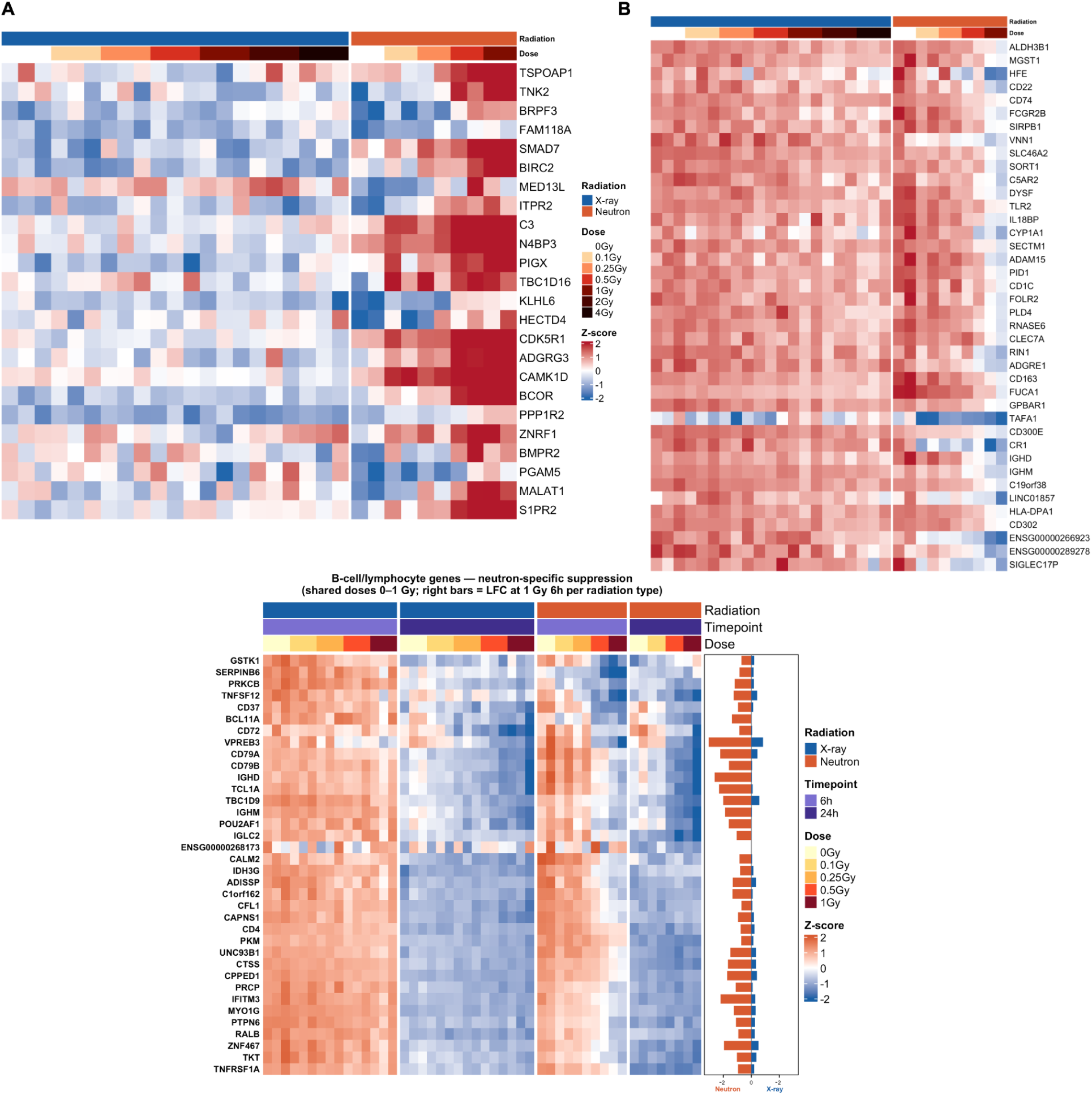
Radiation quality-specific signature. (A) Heatmap of Class A neutron-specific upregulated genes (X-ray-insensitive at 6 h across the full 0–4 Gy dose range). (B) Heatmap of Class A neutron-specific downregulated genes. (C) Heatmap of B-cell and lymphocyte genes showing neutron-specific suppression. Bar plots on the right show radiation type-specific log fold change (LFC) at 1 Gy 6 h for neutrons and X-rays.

The remaining 13 neutron-specifically activated genes were also significantly activated by X-rays at 6 h, but only at higher doses ≥ 2Gy, designated as Class B (dose-shifted). These included the core NF-κB subunits *RELB*, *NFKB1*, *NFKBIA*, *NFKBIB*, *TRAF3*, *OTULIN*, and *TNIP1*, all reaching significance at 4 Gy at 6 h with positive LFCs. This indicated that the NF-κB program is dose-threshold-dependent rather than radiation-quality-specific, with X-rays requiring 2–4 times higher doses compared to neutrons. An exception is *KLF10*, which showed directional divergence between radiation types, potentially reflecting genuinely distinct regulation of this TGF-β/SMAD-responsive transcription factor under high-LET versus low-LET conditions.

### Neutron-preferential suppression: early shutdown of B-cell identity

Out of the DiD-significant genes, 186 unique genes were suppressed preferentially by neutrons within the shared 0-1 Gy dose range. These genes were dominated by B-cell and immune effector components. Similarly, cross-referencing with X-ray 6 h responses across the full dose range yielded a two-class structure. Class A downregulated DEGs comprised 122 genes (66%), whereas Class B downregulated DEGs included 64 genes (34%) that became significant after 2 Gy (36 genes) or 4 Gy (28 genes) X-rays. Thus, the majority of neutron-preferentially suppressed genes showed genuine X-ray insensitivity at the 6 h timepoint across the entire tested dose range. The 122 Class A genes primarily represent components of neutron-specific immune suppression. Among the most strongly suppressed genes after 1 Gy were *IGHD*, *HFE*, *CD163*, *FOLR2*, ENSG00000266923, *CR1*, *FCGR2B*, *RNASE6*, *C5AR2*, and *MGST1*. Four biological patterns emerged from this class: (i) innate myeloid identity (*ADGRE1*, *CD300E*, *PLBD1*, *MARCHF1*, *AIF1*, and *CLEC7A*) (Figure 4B), suggesting that neutrons specifically repressed innate myeloid effectors at doses where X-rays remained transcriptionally silent for these genes; (ii) antigen presentation and B-cell receptor components (*CD22*, *CD74*, *HLA-DPA1*, *HLA-DQB1*, and *HLA-DRB5*); (iii) complement and Fc receptor signaling (*CR1*, *C5AR2*, *FCGR2B*, and *FCER1G*); and (iv) interferon-related genes (*TLR2*, *MX2*, and *IFITM1*), directly corroborating the broad attenuation of IFN-γ pathway activity identified by GSEA (NES −2.01 to −2.17). These findings confirmed that neutron-specific IFN pathway suppression extended from upstream sensing to downstream effector expression.

The Class B genes revealed a dose-shifted pattern of B-cell suppression between radiation qualities. Genes reaching significance 6 h after 2 Gy X-rays included core B cell receptor (BCR) complex components (*CD79A* and *VPREB3*), the signaling adaptor *OSBPL10*, and immune regulators (*CPPED1*, *TBC1D9*, and *NIBAN3*). Genes requiring 4 Gy X-ray at 6 h for significance included *TCL1A*, *POU2AF1*, *BANK1*, *BTK*, *CD79B*, and *MS4A1*, indicating that B-cell suppression reaches significance at lower neutron doses compared to X-rays at 6 h.

The difference in B-cell suppression between radiation qualities is also strongly time-dependent: neutrons reached this endpoint at 6 h while X-rays approached it gradually by 24 h at the same dose (Figure 4C). At 6 h, suppression was exclusive to neutrons across the shared dose range. By 24 h, X-rays induced significant suppression (*TCL1A*, *CD79A*, *IGHD*, *IGHM*, *MS4A1*; all p_adj_ < 10^-12^), while neutron-induced suppression intensified (mean LFC: −1.82 at 6 h vs. −2.52 at 24 h). Notably, even after temporal convergence, neutrons maintained stronger suppression at 24 h (mean LFC: −2.52 for neutrons vs. −1.94 for X-rays).

### Radiation qualities trigger distinct transcriptional kinetics

To examine radiation-quality-specific transcriptional kinetics, a within-radiation-type timepoint DiD analysis comparing baseline-corrected dose responses at 24 h versus 6 h was performed within each radiation type (Supplementary Table 5).

Experimental group-level classification revealed distinct temporal patterns between X-rays and neutrons (Figure 5A). After X-rays, the transcriptional response progressively built up over time. The class of genes showing a response only at 6 h was small and restricted to higher doses (21 genes at 2 Gy and 79 at 4 Gy), including *CD4*, *CD40*, and *CD83*. The dominant class emerged after 24 h across all doses (27 genes at 0.5 Gy, 83 at 1 Gy, 175 at 2 Gy, and 325 at 4 Gy; 610 total), including *MPEG*, *RTL5*, and *NFAM1*. The temporal DiD GO:BP enrichment expanded progressively with X-ray dose (Supplementary Table 6). At 0.5 Gy, enrichment was restricted to B-cell biology, including B-cell receptor signaling, B-cell activation, B-cell differentiation, and humoral immune response, reflecting the initial emergence of B-cell suppression at 24 h (Figure 5B). At 1 Gy, enrichment broadened to include Fc-epsilon receptor signaling, antigen processing and presentation, and lymphocyte proliferation. At the Hallmark level, Allograft Rejection became significant from 1 Gy onwards. At 2 Gy, IFN-γ emerged, while at 4 Gy, TNF-α Signaling via NF-κB, Inflammatory Response, Complement, and IL2-STAT5 Signaling were all significantly enriched (Supplementary Table 7).

**Figure 5:**
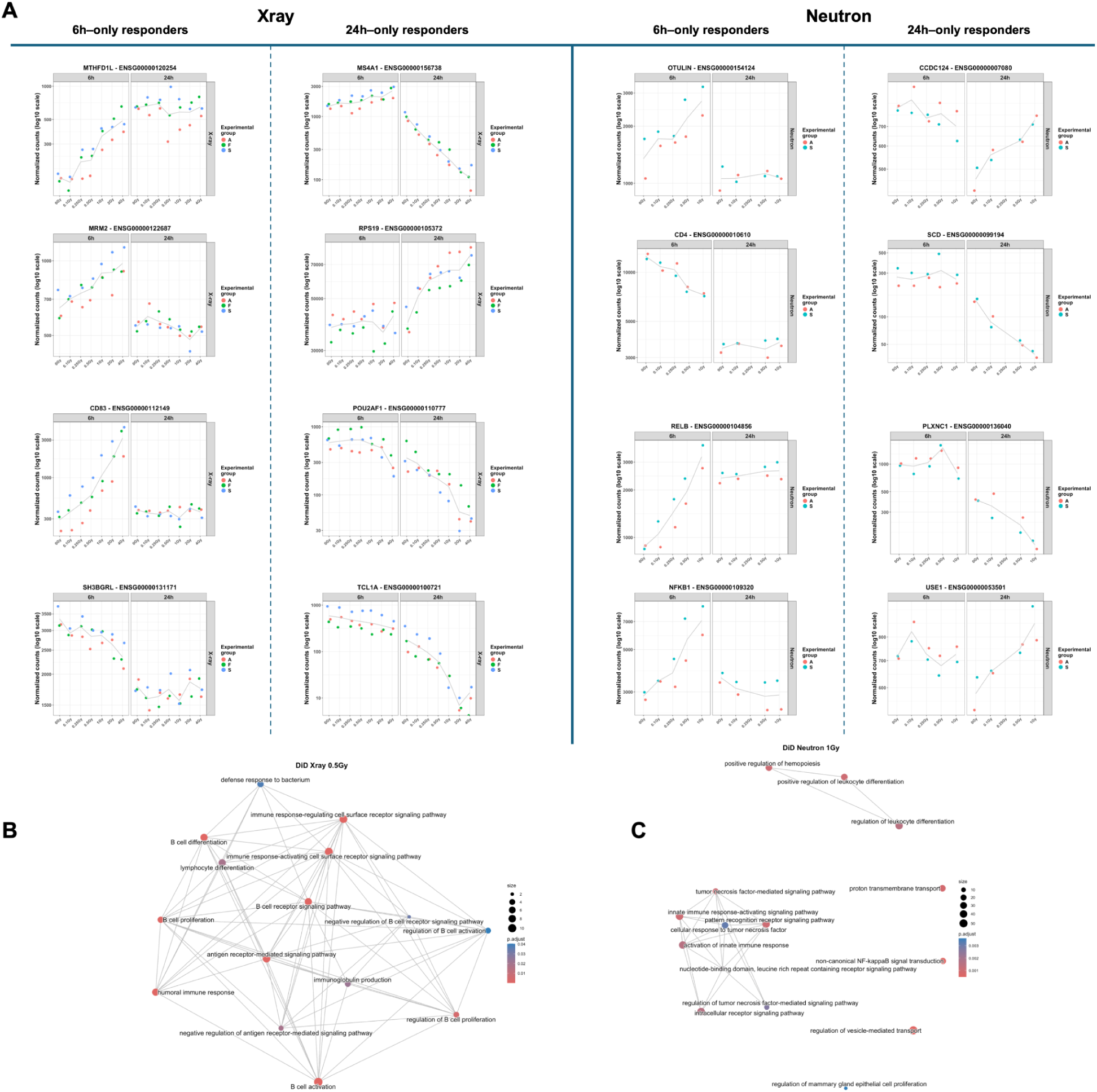
Temporal DiD analysis. (A) Dose-response plots for representative genes showing X-rays 6 h-only responders (left panel, genes significant at 6 h resolving by 24 h), X-rays 24 h-only responders (center-left, genes significantly respond *de novo* at 24 h), neutrons 6 h-only responders (center-right, genes peaking at 6 h and resolving by 24 h), and neutrons 24 h-only responders (right, genes with delayed or sustained neutron responses). Normalised counts on a log10 scale; coloured dots represent individual donors. (B) GO:BP network plot of pathways enriched among DiD-significant temporal genes at 0.5 Gy X-rays. Node size represents gene set size; colour represents adjusted p-value. (C) GO:BP network plot of pathways enriched among DiD-significant temporal genes at 1 Gy neutrons.

Notably, the B-cell receptor signaling enrichment observed 24 h after X-ray exposure, as shown by the temporal DiD analysis, did not represent a *de novo* immune modulation when comparing radiation qualities. Corresponding genes suppressed by X-rays at 24 h were already suppressed by neutrons after 6 h. As these radiation responses converged by 24 h, no radiation-quality-specific DiD effects were detected at that time.

For neutrons, in contrast, the dominant class comprised genes showing a significant radiation response at 6 h (138 genes at 0.5 Gy and 350 at 1 Gy; 488 total) that became non-significant by 24 h. This class was predominantly downregulated (78 down and 60 up at 0.5 Gy; 247 down and 103 up at 1 Gy) and included NF-κB pathway components such as *RELB*, *NFKB1*, and *N4BP3*, whose strong 6 h attenuation diminished over time as *OTULIN*, *NFKBIA*, *NFKBIB*, and *TNIP1* constrained sustained signaling. In contrast, the 24 h-responsive class was smaller (27 down and 19 up at 0.5 Gy and 91 down and 105 up at 1 Gy; 242 total) and included genes such as *SCD*, *RPS19*, *TM7SF3*, and *TAF2*. Significant temporally-enriched pathways were detected only at 1 Gy (Supplementary Table 8). At this dose, 138 significant GO:BP terms reflected attenuation of the acute innate immune response, affecting non-canonical NF-κB signal transduction, tumour necrosis factor-mediated signaling, pattern-recognition receptor signaling, NLR inflammasome signaling, and regulation of leukocyte differentiation (Figure 5C). Three Hallmark pathways were also significant: Apoptosis, Complement, and TNF-α Signaling via NF-κB.

Together, neutrons elicited an early transcriptional response at 6 h, marked by NF-κB activation, upregulation of the complement pathway, and suppression of B-cell identity genes, a signature captured by 488 genes uniquely expressed at 6 h, which largely vanished by 24 h. In contrast, X-rays induced a delayed, progressive response, yielding 611 genes exclusively detected at 24 h. By 24 h post-irradiation, both radiation qualities strongly suppressed B-cell identity genes. This reflects a temporal convergence toward a shared immunosuppressive state, one that neutrons reach rapidly, and X-rays approach gradually. Reversal in the direction of gene expression between 6 h and 24 h was rare (34 genes for neutrons, 12 genes for X-rays). Thus, the impact of radiation quality on the transcriptional response is not merely a matter of magnitude; it is also fundamentally a matter of timing.

### Transcriptomics-based neutron RBE

Finally, as a general measure of neutron effectiveness relative to X-rays, the neutron RBE was determined based on a global dose-dependent gene signature at the whole transcriptome, pathway, and individual-gene levels.

### Global neutron RBE

A composite radiation-responsive gene signature was generated to derive a single summary measure of neutron RBE. Genes with significant dose-dependent expression changes in both radiation types at the same timepoint were identified and ranked by mean absolute slope, yielding a shared directional signature of up to 50 upregulated and 50 downregulated genes per timepoint. Dose-response slopes for the shared gene signature were 1.24 (X-rays, 6 h), 1.61 (neutrons, 6 h), 1.58 (X-rays, 24 h), and 1.91 (neutrons, 24 h). The resulting global neutron RBE was 1.30 (95% CI: 1.14–1.49) at 6 h and 1.21 (95% CI: 1.14–1.28) at 24 h.

### Pathway-level neutron RBE

The pathway-level neutron RBE was computed using GSVA enrichment scores for all GO:BP and MsigDB Hallmark pathways to identify mechanisms underlying the global and gene-level RBE. This analysis identified 76 and 270 pathways with neutron RBE estimates at 6 h and 24 h, respectively (Supplementary Table 9).

At 6 h, 13 activated and 11 suppressed pathways showed neutron RBEs > 1 (Figure 6A), including activation of a cluster of p53-associated DNA damage and apoptosis pathways: regulation of thymocyte apoptotic process (RBE 1.81), thymocyte apoptotic process (RBE 1.37), intrinsic apoptotic signaling by p53 class mediator (RBE 1.28), and DNA damage response signaling by p53 class mediator (RBE 1.19), consistent with the stronger early p53-mediated damage response after neutrons.

**Figure 6:**
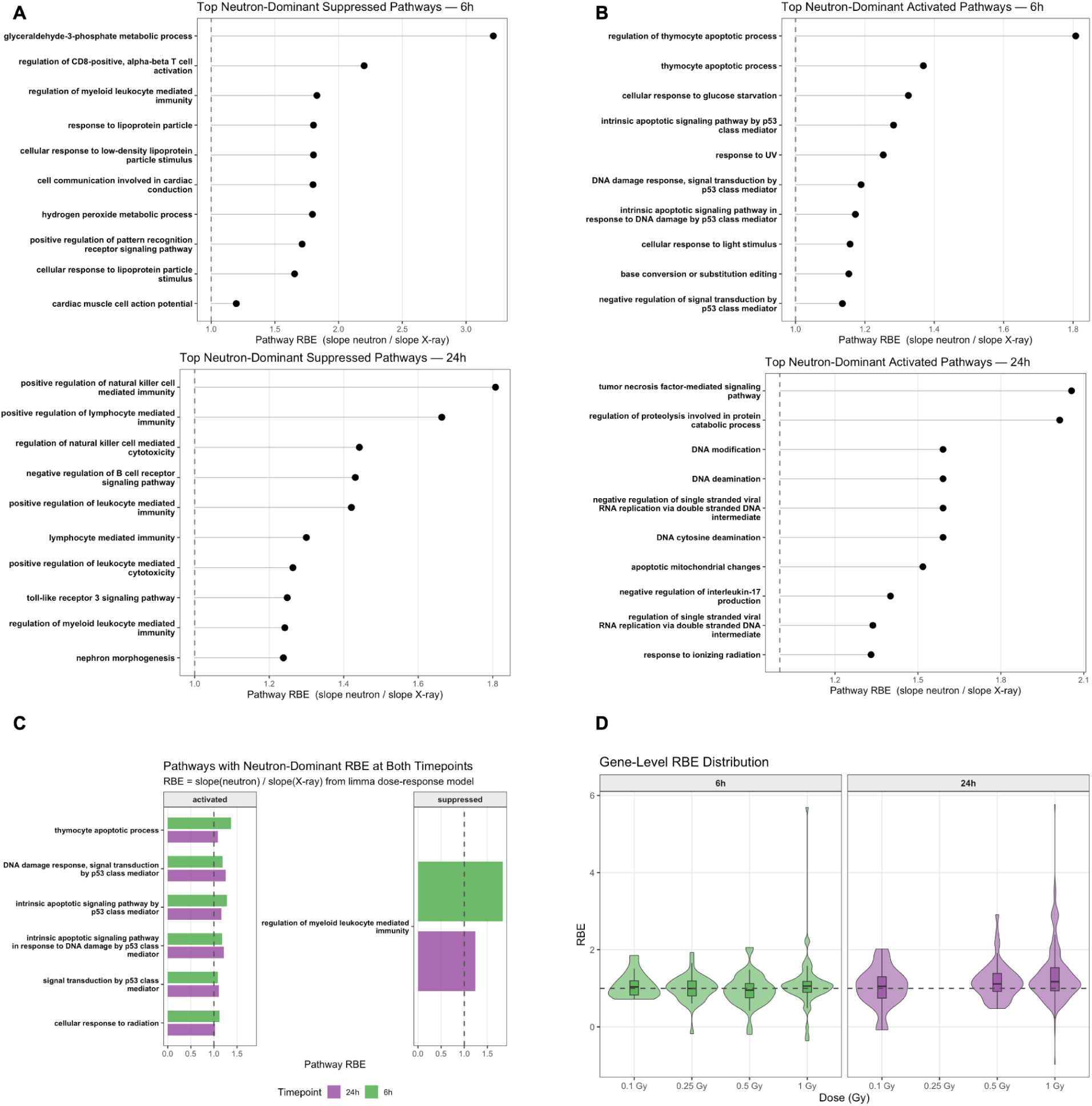
Transcriptomic relative biological effectiveness (RBE) at the pathway and gene level. (A) Dot plot of the top 10 neutron-dominant suppressed pathways at 6 h (upper panel) and 24 h (lower panel). (B) Dot plot of the top 10 neutron-dominant activated pathways at 6 h (upper panel) and 24 h (lower panel). (C) Bar chart of pathways showing consistently neutron-dominant pathways at both 6 h and 24 h post-exposure, split into activated (left) and suppressed (right) panels. (D) Violin plots showing the distribution of gene-level RBE values at 6 h (green, left) and 24 h (purple, right) across all concordant genes at each shared dose.

At 6 h, the suppressed pathway with the highest neutron RBE of 3.21 was glyceraldehyde-3-phosphate metabolic process, consistent with the mitochondrial and glycolytic stress responses described above. High neutron RBEs were also observed in several downregulated immune-related pathways, including regulation of CD8-positive alpha-beta T cell activation (RBE 2.20), regulation of myeloid leukocyte mediated immunity (RBE 1.83), and positive regulation of pattern recognition receptor signalling (RBE 1.71). 24 h after irradiation, neutron-related suppression was the predominant pattern, with 43 downregulated and 20 activated pathways showing a neutron RBE > 1. A cluster of immune surveillance pathways, including NK cell-mediated immunity (RBE 1.81), lymphocyte-mediated immunity (RBE 1.66), natural killer cell cytotoxicity (RBE 1.44), and B cell receptor signaling pathway (RBE 1.43), showed suppression, confirming that neutrons more potently downregulate cellular immune responses in PBMCs at 24 h. Among activated pathways, TNF-mediated signaling (RBE 2.06) and regulation of proteolysis (RBE 2.01) showed the strongest neutron dominance, consistent with enhanced secondary stress and cell death signaling (Figure 6B).

Seven pathways showed neutron RBEs > 1 at both post-exposure timepoints, representing the most reproducible pathway-level findings (Figure 6C). Five were mechanistically linked to p53 and apoptosis pathways: DNA damage response signaling by p53, intrinsic apoptotic signaling in response to DNA damage by p53, intrinsic apoptotic signaling by p53 class mediator, signal transduction by p53 class mediator, and thymocyte apoptotic process, with neutron RBEs ranging from 1.08-1.37 across both timepoints. Regulation of myeloid leukocyte-mediated immunity also showed consistent suppression after neutrons at both timepoints (RBE 1.83 at 6 h and 1.24 at 24 h).

Together, the pathway-level analysis identified two parallel biological mechanisms through which the neutron RBE manifests: a stronger activation of DNA damage sensing and apoptosis centered on p53 signaling, and stronger suppression of immune surveillance pathways including NK cell, lymphocyte, and B cell mediated immunity.

### Gene-level neutron RBE

To compare neutron and X-ray transcriptional responses across genes and dose levels, gene-level neutron RBE was calculated as the ratio of neutron to X-ray LFCs at each dose and time point (Supplementary Table 10). The overall distribution showed a time-dependent increase in neutron RBEs (Figure 6C). At 6 h, 52% of qualifying genes exhibited neutron RBEs > 1, increasing to 67% at 24 h. Median neutron RBEs also increased monotonically with dose (1.05, 1.11, and 1.17 at 0.1, 0.5, and 1 Gy, respectively). The number of genes with neutron RBEs > 2 after 1 Gy increased from 5 genes at 6 h to 41 genes at 24 h, whereas higher X-ray sensitivity (RBE < 0.5) remained rare at both time points (10 and 8 genes, respectively). As already indicated by the temporal DiD analysis, this progressive shift was not driven by the *de novo* emergence of neutron-specific genes at 24 h, but by disproportionately stronger LFCs of shared responsive genes over time (Figure 6D).

Within the shared dose range at 6 h, *CDKN1A* showed consistently elevated neutron RBEs (1.85 at 0.1 Gy, declining to 1.01 at 1 Gy). *ZMAT3* (median RBE 1.93) and *EDA2R* (median RBE 2.12) exhibited the highest consistent neutron RBEs among genes robustly responsive to both radiation types. Additional genes showing elevated neutron sensitivity across doses at 6 h included *MDM2* (3 doses, median RBE 1.25), *PHLDA3* (4 doses, median RBE 1.21), *FDXR* (4 doses, median RBE 1.19), *TP53INP1* (2 doses, median RBE 1.53), and *IER5* (3 doses, median RBE 1.42). In contrast, the nucleotide excision repair genes *XPC* (median RBE 0.94) and *AEN* (median RBE 0.95), the pro-apoptotic effectors *BAX* (median RBE 0.90) and *BBC3* (median RBE 0.99), and the ribosomal p53 regulator *RPS27L* (median RBE 0.81) consistently showed neutron RBEs ≤ 1.0 at 6 h, indicating similar dose-dependent activation of downstream apoptotic and DNA repair effectors by X-rays and neutrons at this timepoint. Together, these findings suggest that the elevated neutron RBE at 6 h is concentrated at upstream damage-sensing nodes rather than downstream effector pathways.

At 24 h, the highest gene-level neutron RBEs were observed predominantly among B-cell- and lymphocyte-related genes. *LMO2* (median RBE 2.93), a transcription factor essential for hematopoietic progenitor and B-cell development, together with *POU2AF1* (median RBE 1.80), *CD180* (median RBE 1.96), and *SCIMP* (RBE 3.01), showed sustained and deepened suppression after neutrons at 24 h. Immunoglobulin chain genes *IGKC* (RBE 3.49) and *IGKV3-20* (RBE 2.97) were also among the most neutron-sensitive genes at 24 h, further supporting stronger disruption of the B-cell transcriptional program by neutrons. These findings directly connect the gene-level neutron RBE framework to the B-cell identity suppression program identified by the DiD analysis, providing independent quantitative evidence that neutron irradiation suppresses B-cell identity not only earlier, but also more deeply and persistently. The long non-coding RNA *LINC01857* showed the highest individual neutron RBE of 5.77 after 1 Gy at 24 h.

Notably, two genes exhibited temporal neutron RBE reversals between timepoints. *BAX* and *CMBL* exhibit a time-dependent shift in radiation sensitivity, with higher X-ray sensitivity at 6 h (*BAX* median RBE 0.84; *CMBL* median RBE 0.70) and increased neutron responsiveness at 24 h (*BAX* median RBE 1.19; *CMBL* median RBE 2.5).

Integration of the three analytical approaches for neutron RBE assessment, by global slope ratios, gene-level LFC ratios, and pathway-level GSVA slope ratios, provided concordant evidence for stronger biological effects of neutrons across multiple levels of biological organization.

## Discussion

Neutrons play a critical role in many planned and accidental radiation exposure scenarios. However, the transcriptional response to neutrons and high-LET particle radiation in general, which is increasingly assessed for biodosimetry in peripheral blood samples, has not yet been systematically characterized, either in this specific context or with respect to broader molecular and immunological responses and associated health effects. A particular challenge remains the discrimination between different radiation qualities in exposure scenarios involving mixed radiation fields. We therefore conducted a comprehensive genome-wide characterization of transcriptional responses in PBMCs isolated from *ex vivo* irradiated human whole blood following exposure to an accelerator-derived fission-like spectrum of neutrons compared with X-rays at 6 and 24 h post-exposure.

The transcriptional variance was dominated by time post-exposure, followed by radiation quality and inter-donor differences. These factors were accounted for in the analyses to extract gene sets that are specific to time, dose, and radiation quality while minimizing the influence of inter-donor variability.

Due to its LET-dependent DNA-damaging effects, ionizing radiation induces a coordinated DNA damage response predominantly mediated by p53-centered signaling pathways. We confirmed this at the transcriptional level for all doses, time points, and both radiation qualities, as demonstrated by the dose-dependent up-regulation of the p53 downstream targets *FDXR*, *BBC3*, *GADD45A*, *DDB2*, and *CDKN1A*. This finding aligns with the extensive biodosimetry literature that has validated these genes in blood samples across a wide range of radiation-exposure scenarios (13,33–38). This study identified the seven genes *GADD45A*, *PCNA*, *XPC*, *FDXR*, *TRIAP1*, *AEN*, and *ZNF79* as the highest-confidence radiation-quality-independent biodosimetry candidates in a formally controlled human *ex vivo* study across multiple experimental settings, exhibiting perfect dose-response correlations across all conditions with donor SD = 0. Additionally, *PVT1*, *TMEM30A*, *ISG20*, *ASTN2*, *MAP4K4*, *ACTA2*, and *EDA2R* showed near-perfect dose-response correlations with minimal inter-donor variability (SD < 0.1). *PRKAB1* (AMPK β1 subunit) is a less-studied radiation response gene, but its SD of 0.014 across all experimental conditions warrants further functional investigation.

Previously, we also identified a set of hitherto undescribed radiosensitive genes in *ex vivo* X-irradiated human blood (12), several of which were recapitulated and validated in the present study. Four upregulated genes (*MIR34AHG*, *PMAIP1*, *MRM2*, and *TIFA*) and five downregulated genes (*NIBAN3*, *CORO1A*, *IFIT3*, *PTPRS*, and *RBBP8*) were differentially expressed, whereas neutron irradiation induced differential expression of three upregulated genes (*PMAIP1*, *MIR34AHG*, and *TIFA*) and two downregulated genes (*PLEKHG1* and *NIBAN3*). A certain inconsistency and fluctuation in gene signatures can be expected and explained by variations in technical, experimental, or analytical settings.

In general, neutrons elicited an earlier and stronger transcriptional response at the same dose levels compared with X-rays, consistent with the greater biological impact of high-LET radiation. This is particularly evident in the increased p53-downstream signaling by higher complexity of DNA damage. Fast neutrons induce clustered and complex DNA damage through indirect ionization processes mediated by energy-dependent generation of secondary high-LET particles, thereby increasing RBE and associated health risks. Maximum neutron RBEs can reach 20 depending on the endpoint and energy (typically 0.3-0.4 MeV), and are usually inversely correlated with neutron dose (1).

At a dose of 1 Gy, neutrons induced a 12-fold higher number of DEGs at 6 h compared with X-rays. Although this number decreased by 24 h, the initially triggered transcriptomic response deepened, whereas X-rays showed an increase in DEGs over the same period. This divergent temporal trajectory, an early transcriptomic burst followed by consolidation for neutrons, versus a gradual expansion for X-rays, likely reflects distinct damage-sensing kinetics between radiation types, with high-LET clustered DSBs driving a rapid but partially self-resolving acute response. Despite these differences in magnitude and dynamics, both radiation qualities activated a p53-mediated DNA damage response even at low doses; neutrons, however, additionally engaged immunogenic signalling at doses where X-rays elicited only a p53-centred genotoxic stress response.

A more immunogenic response can also be triggered by clustered DNA damage induced by high-LET radiation. Besides complex chromosomal rearrangements, this leads to increased formation of cytoplasmic DNA fragments that activate the cGAS-STING pathway at lower doses than low-LET photon irradiation (39). This mechanism may explain the immunogenic responses observed after neutron exposure at lower doses compared with X-rays, as well as the dose-threshold architecture of the DiD findings. Specifically, the cGAS-STING-NF-κB program was engaged at 1 Gy for neutrons, whereas X-rays required doses of ≥ 2 Gy, indicating a quantitative rather than qualitative difference in innate DNA-damage-associated immunosensing efficiency per unit dose.

One of our primary objectives was to develop a previously unavailable concept for a transcriptome-wide neutron RBE, aimed at identifying specific patterns or biomarkers that distinguish the biological responses between radiation qualities. To our knowledge, this study provides the first formal quantification of a time-dependent genome-wide transcriptomic neutron RBE in human PBMCs isolated from irradiated whole blood, computed across three analytical levels: (i) global slope ratio, (ii) gene-level fold-change ratio, and (iii) pathway-level GSVA slope ratio. The strong concordance across all three levels in assessing neutron transcriptomic RBE provides a robust and biologically meaningful endpoint that may serve as a novel transcriptomic calibration metric for neutron and mixed-field biodosimetry.

Previous studies estimating neutron RBEs based on gene expression in human blood have relied primarily on microarray platforms or qRT-PCR analyses focused on predefined sets of radiation-responsive genes (13,17,19). Broustas et al. (13) investigated the transcriptional response to an accelerator-generated broad-energy neutron spectrum (0.2–9 MeV), delivered at doses of 0.1, 0.3, 0.5, or 1 Gy, compared to X-rays, 24 h post-irradiation. Confirmation of whole-genome microarray data for 8 selected neutron-dominated or non-dominated genes among 125 radiation responders by qRT-PCR showed median neutron RBE values of 0.8–1.2 for *BAX*, *TNFRSF10B*, *ITLN2*, and *AEN*. In contrast, *VWCE*, *FDXR*, *PHLDA3*, and *EDA2R* had higher median neutron RBEs (1.25–3.65), with a slight increase at higher doses. The overall neutron RBE across all these genes was approximately 1.8.

More recently, Palmqvist et al. (19) employed the same neutron field as used in this study to examine the expression of six radiation-responsive genes in irradiated blood samples from three healthy donors. Gene expression was quantified 4 h post-irradiation at doses of 0.5, 0.75, and 1 Gy using qRT-PCR, and compared to responses induced by gamma-rays. The average neutron RBE values, derived from linear dose-response regressions, were also both dose- and gene-dependent, ranging from 1.39 to 3.91. Specifically, the mean RBEs were: *FDXR* = 1.39, *BBC3* = 1.80, *XPC* = 1.75, *MDM2* = 2.08, *CDKN1A* = 3.59, and *GADD45A* = 3.91. Together, the data from previous studies, consistent with our own, demonstrate that transcriptomic neutron RBE values are both gene- and dose-dependent, spanning a range of 1 to 4, with the majority falling within the range of 1–2.

Of note, Palmqvist et al. (19) also investigated various sequential irradiation schemes, including neutrons followed by photons, and *vice versa*, and found purely additive effects of both radiation qualities. This observation underscores the value of our transcriptomic RBE data for individual radiation qualities also in the context of mixed-field exposures, where accounting for the contribution of each component is essential for accurate biodosimetry and risk assessment.

However, our first analysis of two post-exposure time points revealed a slightly lower neutron RBE at 24 h compared to 6 h. This suggests that timepoint-specific corrections should be applied when using transcriptomic neutron RBE values across different post-irradiation timepoints. At the level of individual genes, a gene module with high neutron RBEs could be identified based on a dose-dependent increase in neutron RBE observed at 24 h, where 14.1% of concordant genes exhibit a neutron RBE > 2 at 1 Gy, compared to 5.3% at 0.1 Gy. Its preferential amplification under high-LET conditions may provide an independent 24 h quality signal driven more by effect magnitude than by the mere presence or absence of expression.

The neutron potency was not uniformly distributed across the p53 programme but concentrated at upstream damage-sensing and amplification nodes: *CDKN1A*, *FDXR*, *ZMAT3*, and *EDA2R* all showed consistent neutron-dominant RBEs across multiple doses, while downstream apoptotic effectors (*BAX*, *BBC3*, *XPC*, *AEN*) showed neutron RBE values near 1.0 at 6 h, and *BAX* itself reversed from X-ray-dominant at 6 h to neutron-dominant at 24 h. This upstream-weighted p53 response occurred alongside a parallel, neutron-specific innate-immune signature centered on canonical and non-canonical NF-κB (*RELB*, *NFKB1*, *NFKB2*), cGAS-STING amplifiers (*PGAM5*, *TSPOAP1*, *ITPR2*, *N4BP3*), and complement C3, a coordinated response absent from all X-ray 6 h samples across the shared dose range (0–1 Gy). DNA damage signaling and NF-κB activation are known to be co-induced downstream of ionizing radiation (40–42), consistent with the concurrent engagement of both programmes observed here under neutron but not X-ray exposure. Non-canonical NF-κB, which operates via NIK-IKKα-p100/p52, is the primary NF-κB pathway governing B-cell homeostasis and dendritic cell maturation (43); its early activation by neutron-induced cGAS-STING at 1 Gy provides a plausible mechanistic basis for the broader and deeper B-cell suppression observed at that dose.

B-cell suppression may also be reinforced epigenetically through BCOR-mediated histone H2A ubiquitination, leading to silencing of B-cell loci, alongside S1PR2-driven apoptotic signalling (44–46). The concurrent induction of C3 with suppression of complement receptors *CR1* and *C5AR2*, myeloid pattern recognition receptors (*CLEC7A*/Dectin-1, *TLR2*, *ADGRE1*), and IFN-γ response pathways (NES −2.01 to −2.17) supports a broader early immunosuppressive phenotype, consistent with previous neutron transcriptomics data in mice (15,16). The strong neutron-indicator *MALAT1*, identified as the most strongly induced Class A gene, is also transcriptionally regulated by NF-κB and additionally by p53 via distinct binding sites in its proximal coding region (47), accounting for its characteristic neutron-specific response. Notably, low plasma *MALAT1* levels have been linked to radiotherapy-related adverse effects and poor prognosis in cancer patients (48).

While the majority of transcriptomic biodosimetry studies have focused on identifying radiation-quality-independent responders, we additionally identified a set of genes specifically regulated by neutron irradiation, providing a candidate signature for radiation quality discrimination. This Class A signature spans orthogonal yet mechanistically coherent pathways, making it unlikely to be mimicked by radiation-unrelated inflammatory confounders that activate NF-κB without simultaneously suppressing BCR and MHC class II signaling. Beyond *MALAT1*, the strongest Class A genes included *C3*, *RELB*, *NFKB1*, *KCNH4*, and *MIAT* (upregulated) and *IGHD*, *VPREB3*, *TCL1A*, *CD163*, *CR1*, *FCGR2B*, *CLEC7A*, and *TLR2* (downregulated), enabling construction of a predictive gene set score based on their opposing regulation. For example, a composite Class A gene set score integrating NF-κB activation (e.g., *RELB*, *NFKB1*, *C3*) with B-cell/myeloid suppression (e.g., *IGHD*, *CLEC7A*, *CD79A*) could enhance discrimination of radiation quality beyond any single gene. This signature shows no X-ray response even at 4 Gy 6 h; however, their activation threshold may lie beyond the maximum X-ray dose tested, or they may respond only at later timepoints following X-ray exposure.

In addition to the Class A signature, *ZMAT3* and *EDA2R*, which showed the highest consistent RBE among concordant genes (median RBE 1.93 and 2.12, respectively), could serve a dual biodosimetric role as universal dose markers already included in validated panels and as candidate markers of radiation quality by comparison with photon-calibrated reference curves (37,49).

The 0.5 Gy dose threshold for DiD significance radiation qualities suggests that this dose represents a threshold for radiation-type discrimination. Furthermore, the opposing kinetic profiles within each radiation type, 405 unique genes peaking at 6 h after neutron exposure and resolving by 24 h, versus 396 unique genes emerging only at 24 h within the X-ray response, constitute a potential transcriptional clock. The ratio between resolving and emerging expression patterns may enable estimation of exposure timing independently of dose. Temporal transcriptomics for exposure-time assessment, distinct from dose estimation, remains largely unexplored in biodosimetry and warrants further investigation.

Beyond radiation accident scenarios involving neutrons and biodosimetry, our findings are relevant to the role of neutrons, often secondarily generated by interactions of primary radiation with target materials, in radiotherapy (using very high-energy photons, particle beams, or boron neutron capture therapy), occupational exposures in nuclear environments, and manned spaceflight (1,2,50,51). In these settings, our data provide insights into health consequences beyond the well-established heightened DNA-damaging and mutagenic potential of neutrons, particularly regarding immunomodulation and immunosuppressive effects.

One limitation of our study was the small number of donors, owing to the complex logistics of blood donation and exposure at the neutron facility. Still, we covered a relevant dose range and two analysis time points, building on our previous experience with *ex vivo* and *in vivo* transcriptomic human blood photon data (12,52–54).

## Conclusion

This study provides the first comprehensive genome-wide characterization of human PBMCs transcriptional responses to an accelerator-derived fission-like spectrum of neutrons in comparison with photon radiation, revealing distinct time- and dose-dependent molecular response patterns. We identified both a radiation quality-specific gene signature with strong discriminatory potential and a universal biomarker signature that is robust across donors and highly correlated with radiation dose. Furthermore, we present the first genome-wide, three-layer estimation of the transcriptomic RBE at the gene, pathway, and global transcriptomic levels, providing a comprehensive framework for quantifying radiation quality effects. Together, these findings advance the mechanistic understanding of how radiation quality shapes biological responses and lay the foundation for the development of next-generation transcriptomic biodosimetry approaches capable of both accurate dose estimation and reliable discrimination of radiation quality.

## Supporting information

Supplements_Salah_et_al.

## Availability of code, data, and materials

All analyses presented in this manuscript were run in R v4.5.1, with a detailed computational workflow available at (https://github.com/AhmedSAHassan/Phybion_Neutron_Xray). The processed data together with the metadata of the corresponding samples and the analysis code for reproducibility are available on this GitHub repository.

## Ethics declarations

The studies involving human participants were reviewed and approved by the Ethics committee of the Rhineland-Palatinate Chamber of Physicians [No. 2023-17191].

## Acknowledgements

The authors would like to thank B. Lutz and A. Heiske for their assistance during the neutron irradiations, and O. Döhr, T. Heldt, E. Holland, and J. Pieper for operating the ion accelerators at PTB.

