## Supplementary figures and images for "Comparative Transcriptional Responses of Human Blood to Neutron and Photon Irradiation"

### Supplementary_figure1.png

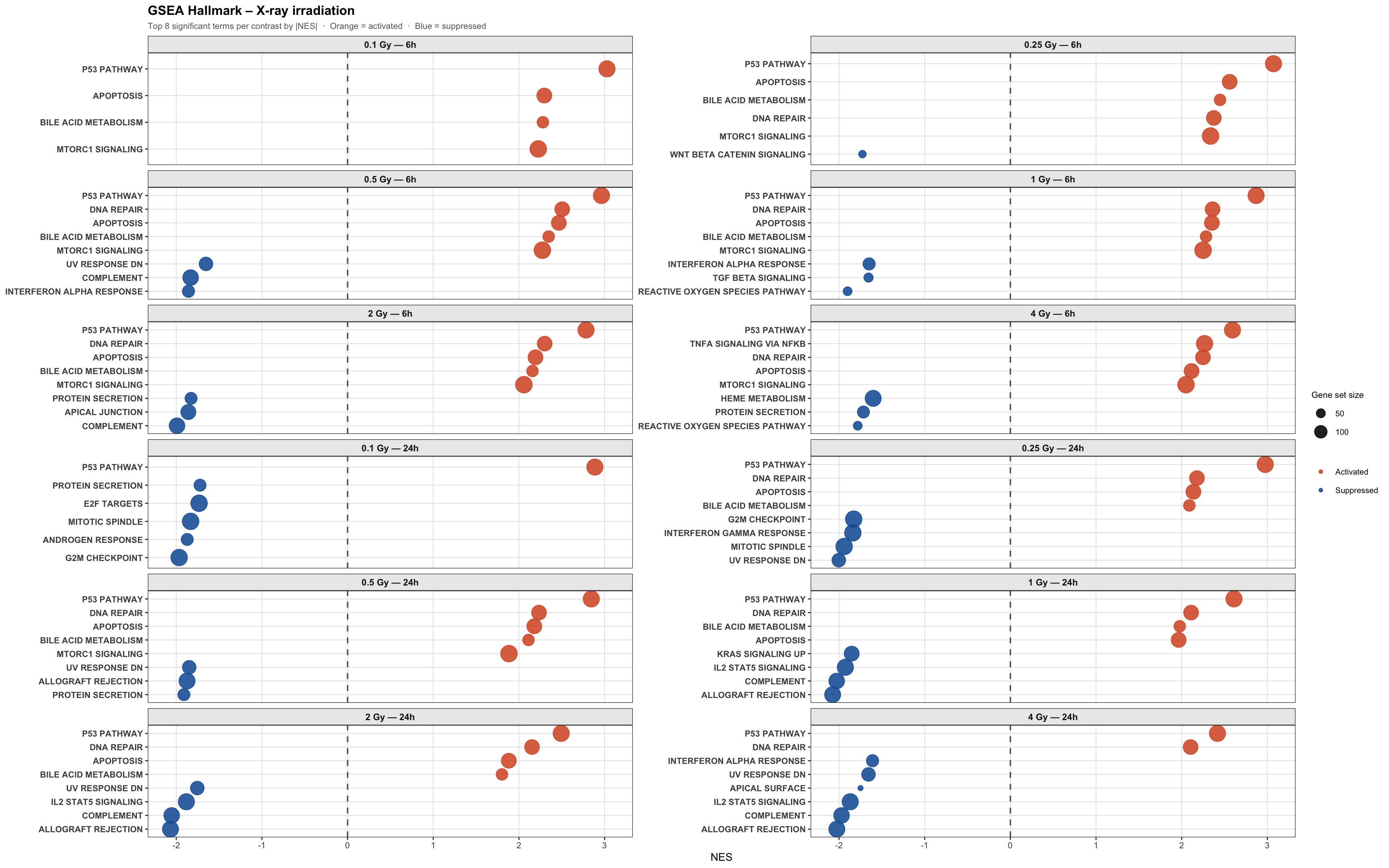
